# ITHANET: Information and database community portal for haemoglobinopathies

**DOI:** 10.1101/209361

**Authors:** Petros Kountouris, Coralea Stephanou, Celeste Bento, Pavlos Fanis, Jacques Elion, Raj S Ramesar, Bin Alwi Zilfalil, Helen M Robinson, Joanne Traeger-Synodinos, Human Variome Project Global Globin 2020 Challenge, Carsten W Lederer, Marina Kleanthous

## Abstract

Haemoglobinopathies are the commonest monogenic diseases, with millions of carriers and patients worldwide. Online resources for haemoglobinopathies are largely divided into specialised sites catering for patients, researchers and clinicians separately. However, the severity, ubiquity and surprising genetic complexity of the haemoglobinopathies call for an integrated website to serve as a free and comprehensive repository and tool for patients, scientists and health professionals alike. This paper presents the ITHANET community portal, an expanding resource for clinicians and researchers dealing with haemoglobinopathies. It integrates information on news, events, publications, clinical trials and haemoglobinopathy-related organisations and experts and, most importantly, databases of variations, epidemiology and diagnostic and clinical data. Specifically, ITHANET provides annotation for 2690 haemoglobinopathy-related variations, epidemiological data for more than 180 countries and information for more than 600 HPLC diagnostic reports. The ITHANET portal accepts and incorporates contributions to its content by local experts from any country in the world and is freely available for the public at http://www.ithanet.eu.

## INTRODUCTION

Haemoglobinopathies are the commonest monogenic diseases in the world, posing a major public health problem worldwide. It is estimated that around 5.2% of the world’s population carry a potentially pathogenic globin gene and that, annually, over 330 thousand newborns bear a serious haemoglobinopathy [1]. Haemoglobinopathies comprise the thalassaemias, the sickle-cell disorders (SCD), haemoglobin (Hb) E, Hb C and other, rarer disorders and are mainly caused by mutations in the two globin-gene clusters, namely the α-globin locus (Chromosome: 16, Accession: NG_000006) and the β-globin locus (Chromosome: 11, Accession: NG_000007), which can cause defects in the structure of Hb or reduced synthesis of globin chains and of Hb within the red blood cells. To date, more than 2000 disease-causing mutations have been reported, affecting different levels of gene regulation and expression [2].

Through resistance of carriers to the malaria parasite, haemoglobinopathies have a high prevalence in former malaria regions in the Mediterranean, the Middle East, South-East Asia and Sub-Saharan Africa [3–5]. However, demographic events, such as migration from high-prevalence areas and the consequent intermixing of populations, have contributed to the spread of haemoglobinopathies worldwide [6–8] and, consequently, their prevalence is rising in non-endemic regions, such as Northern and Western Europe and North America, posing a major challenge for researchers and health professionals. In addition, the lack of high-quality epidemiological data, the high variation of disease prevalence in different ethnic groups and in different regions within the same country, as well as general underestimates of haemoglobinopathy incidence, particularly in low-and middle-income countries, are serious impediments to appropriate policy making [9,10].

Recent advances in biotechnology, particularly the emergence of high-throughput sequencing technologies, as well as recent increased research interest in the field of rare diseases, which affect fewer that one in 2000 population-wide, have led to an explosion in the amount of genetic, clinical and diagnostic information. In the field of haemoglobinopathies, the commonest rare disease in the world, online resources are largely divided into specialised sites catering for patients, researchers and clinicians separately, while available information has been fragmented into different databases, mainly variation-centric and genotype-phenotype databases. More specifically, relevant information can be found in genome-wide resources, such as dbSNP [11], OMIM [12], ClinVar [13] and HGMD [14], but the level of detailed annotation varies, particularly for rare, yet clinically significant, variations [15]. In addition, locus-specific databases, such as HbVar [16] and several databases based on the Leiden Open Variation Database (LOVD) software [17,18], have been successful in the past, but their curation has been a challenging task. Therefore, there is an urgent need for an integrated resource that will bring together all available haemoglobinopathy-related information and will set up a community-based framework for the continuous and consistent collection and integration of high-quality data.

Herein, we present the ITHANET community portal, an expanding resource for clinicians and researchers dealing with haemoglobinopathies, that (a) integrates the latest haemoglobinopathy-related information, including news, events, publications and clinical trials, (b) provides information about the worldwide community of organisations and experts working on haemoglobinopathies, and (c) develops, and maintains databases of variations, epidemiological, diagnostic and clinical data. The ITHANET portal was initially the result of a multinational Euro-Mediterranean project [19] and has progressed to becoming an expanding community resource in collaboration with the Global Globin 2020 (GG2020) Challenge by the Human Variome Project (HVP). GG2020 is now officially linked to the ITHANET portal after review of all known existing databases in the field [20] and with additional partnership through a shared ITHANET-GG2020 Expert Panel application for haemoglobinopathy-related variant classification under the Clinical Genome (ClinGen) Resource. The ITHANET portal welcomes contributions to its content and acknowledges all contributions in a specifically designed section. ITHANET is free and available at http://www.ithanet.eu.

## METHODS

### Data Structure and Management

The ITHANET portal is freely accessible online for viewing, searching and administrating as a website in the form of HTML documents. The application is written in PHP (http://www.php.net) based on the “Joomla!” content management system (http://www.joomla.org) and uses the jQuery JavaScript library (http://www.jquery.com), as well as packages jQuery-UI (http://www.jqueryui.com), DataTables (http://www.datatables.net), HighCharts and HighMaps (http://www.highcharts.com), to enhance the presentation of the data. The interface does not require the installation of additional plugins, such as Flash and Microsoft Silverlight, and, thus, works natively across all modern web browsers and the majority of mobile web browsers. All data available in ITHANET are stored and organised in a relational database using MySQL (http://www.mysql.com), an open-source relational database management system widely utilised in database design in bioinformatics and biomedical informatics. ITHANET is hosted by the Cyprus Institute of Neurology and Genetics (http://www.cing.ac.cy) using Apache 2 HTTP Server (http://www.apache.org).

### Data Collection and Database Curation

A key component for the creation of a public knowledge base is the efficient collection, validation and annotation of relevant information. ITHANET uses a combination of automatic and manual curation to update and enrich the portal’s content. Weekly updates on scientific literature are automatically received from PubMed using the following search query: “*thalassemia [tiab] OR thalassaemia [tiab] OR hemoglobin [tiab] OR haemoglobin [tiab] OR sickle-cell [tiab] OR hemoglobinopathies [tiab] OR haemoglobinopathies [tiab]*”. Subsequently, the references are manually filtered to find relevant information. Importantly, ITHANET is an active partner in and beneficiary of primary research data from several projects on haemoglobin disorders, while additional information is retrieved through subscriptions to scientific societies and organisations. In addition, ITHANET accepts and incorporates contributions to its content by local experts from any country in the world and acknowledges their contribution eponymously in affected content pages of databases IthaGenes and IthaMaps and on a specifically designed page (“Contributors and Curators”).

## RESULTS

As illustrated in Figure 1, existing content on the ITHANET portal can be divided into four main categories: (a) provision of the latest information related to haemoglobinopathies (e.g. news, events, publications, clinical trials), (b) worldwide collection of organisations and experts working on haemoglobinopathies, and (c) databases and computational tools.

**Figure 1:**
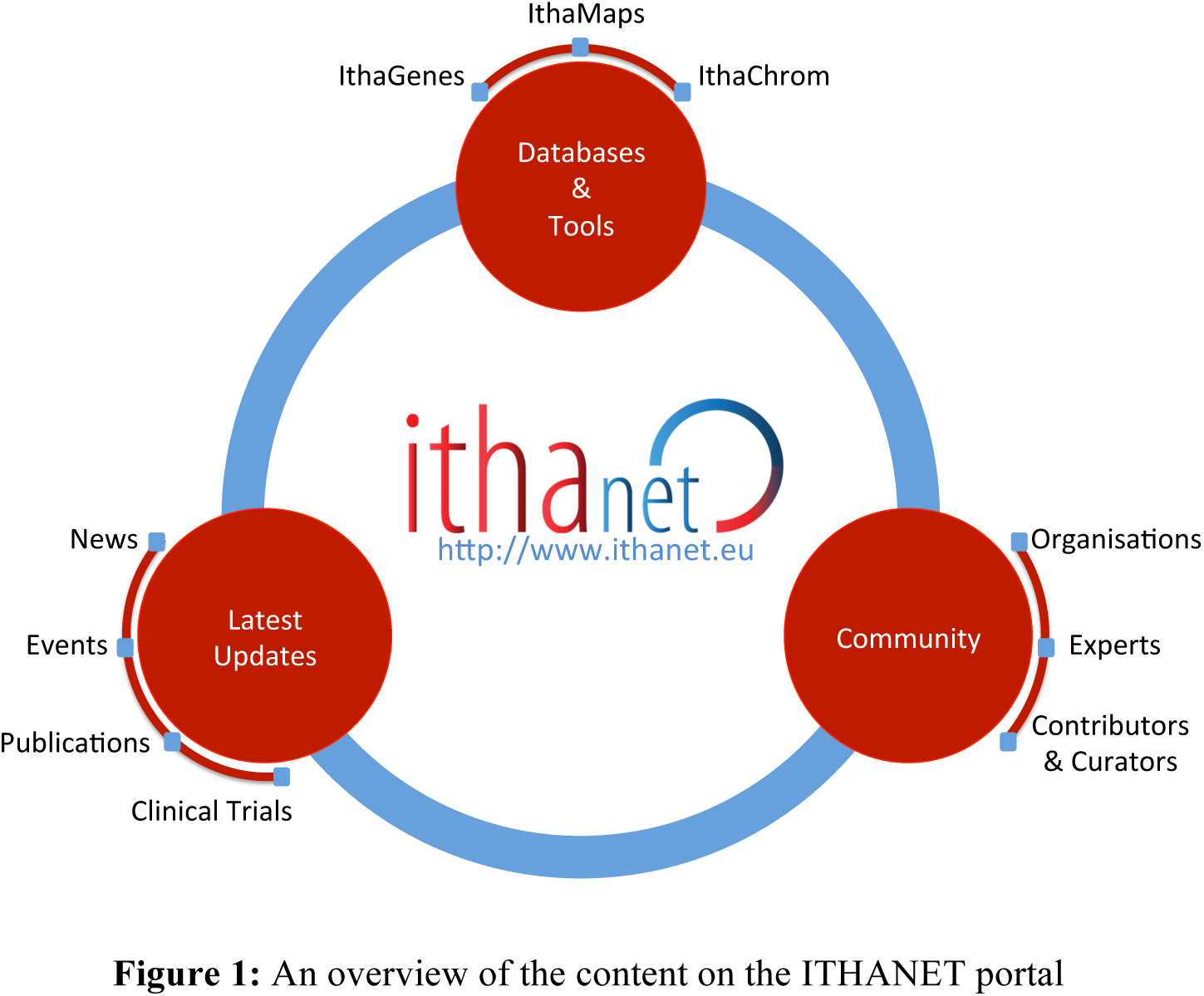
An overview of the content on the ITHANET portal

### Latest Updates

Through its role as a community resource, the ITHANET portal collects and provides the most recent information about haemoglobinopathies and closely related topics from the fields such as human genetics and genomics. Specifically, recent news, such as new research grants, significant discoveries, press releases and clinical trials, are provided in the “News” section, while searchable lists of past and future scientific events, recent publications and clinical trials are provided in sections “Events”, “Publications” and “Clinical Trials”, respectively. Information on scientific events is retrieved through subscriptions to numerous scientific organisations/societies, patient organisations and research networks, while information on publications and clinical trials is retrieved on a weekly basis from NCBI PubMed and clinicaltrials.gov, respectively. In October 2017, manually curated entries of these ITHANET sections comprised 149 news items, 271 past or future events, 3993 publications and 411 clinical trials. Moreover, registered subscribers are informed about the ITHANET updates through a regular electronic newsletter.

### Community

International collaboration and synergies are critical for tackling the burden of rare diseases, in which the number of patients and, consequently, experts specialising on specific diseases is small. In an effort to map institutions and scientists with expertise in haemoglobinopathies and to facilitate collaboration, ITHANET provides a searchable list of organisations related to haemoglobinopathies, with detailed description, contact details and affiliated experts and, similarly, a searchable list of experts working on different areas of haemoglobinopathies with detailed information about contact details, research interests and affiliations and, importantly, possible contributions to ITHANET content. In October 2017, ITHANET stored information about 147 organisations and 174 experts, of whom 29 have also directly contributed to its content in all major sections of the portal, i.e. IthaGenes, IthaMaps and IthaChrom. Such contributions are critical to the accuracy, expansion and growing success of the portal and are therefore acknowledged in a specific section (“Contributors and Curators”).

### Databases and Tools

#### IthaGenes

IthaGenes is a database that organises genes and variations affecting haemoglobinopathies, including disease-causing and disease-modifying variations, as well as diagnostically relevant neutral polymorphisms [2]. IthaGenes entries started out with globin gene causative mutations and annotations initially collated in the books “A Syllabus of Human Hemoglobin Variants (Second Edition)” [21] and “A Syllabus of Thalassemia Mutations” [22] and has since comprehensively incorporated any subsequently reported variations and annotations from recent articles and reports [23–26] and from existing information on other public databases, such as HbVar [16], dbSNP [11], ClinVar [13], OMIM [12] and SwissVar [27]. IthaGenes has six main sections covering the catalogue of genes and regions, the catalogue of mutations, advanced search, statistics, the catalogue of references, and, finally, frequently asked questions (FAQs). Advanced search provides a powerful interface for queries and access to all IthaGenes database entries, with a wide range of search options as previously described [2]. Moreover, IthaGenes integrates the NCBI sequence viewer for detailed graphical representation of each variation and provides phenotype, epidemiology and HPLC data, related publications and external links. Comprehensive coverage, search functionality, extensive annotation and visualisation, systematic inclusion of external links and interconnectedness internally and with other ITHANET portal sections make IthaGenes an exceptional resource for professionals in the field.

Over the past few years, IthaGenes has been established as the largest database of variations related to haemoglobinopathies, with the annotation of around 2000 disease-causing or potentially disease-causing variations. Notably, a joint ITHANET-GG2020 application is currently under review by ClinGen for the assignment of an Expert Panel status towards comprehensive annotation of all variants affecting haemoglobinopathies, based on the guidelines of the American College of Medical Genetics and Genomics (ACMG), and their submission to the NCBI ClinVar database [13]. An ongoing pilot study evaluating the Panel’s haemoglobinopathy-specific rules for a subset of variations will help fine-tune the classification criteria and represents the final step required for Panel approval by ClinGen. The Panel will begin the variant annotation with the genes located in the globin gene clusters, which carry the commonest and most highly penetrant variations, and will subsequently proceed with annotation of variations located in other genes that have a modifying effect on haemoglobinopathies.

The first IthaGenes release [2] was mainly focused on the annotation of variations on the globin gene clusters with a small number of additional modifier genes. However, the enormous and still expanding accumulation of data from genome-wide association studies aiming to identify novel modifiers of haemoglobinopathies has resulted in the accumulation of new genes and loci associated with haemoglobinopathy-related phenotypes. In fact, information on *trans*-acting modifiers is already considered as an important factor in the clinical management of patients with haemoglobinopathies [28]. Thus and after incorporating a large number of genome-wide or targeted association studies over the past two years, the number of genes and loci annotated in IthaGenes has increased from 32 to a total of 232 and, consequently, the number of annotated variations from 1963 to a total of 2690, comprising around 600 disease-modifying variants. In addition, most variants are assigned at least one of 41 clinical phenotypes included in IthaGenes that are now linked with established phenotypic annotations by the Human Phenotype Ontology [29,30] and OMIM [12]. Figure 2 shows the distribution of disease-modifying mutations by their associated phenotype.

**Figure 2:**
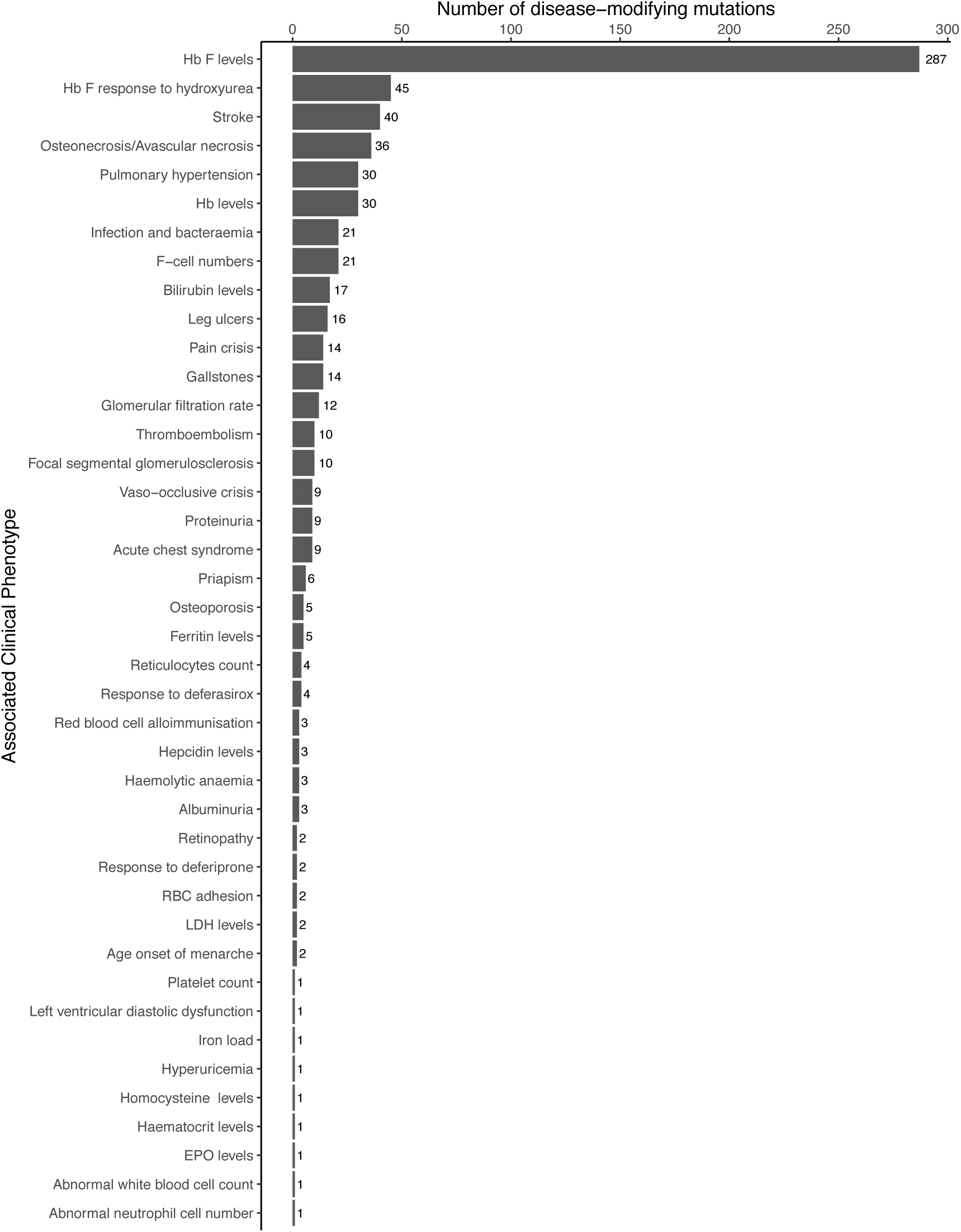
The distribution of disease-modifying mutations by associated clinical phenotype. Many clinical genotypes are interrelated and can be inferred from reported findings, but only explicitly reported phenotypes are included in IthaGenes.

New variations can be directly reported by completing the IthaGenes submission form or by using the ITHANET contact form (http://www.ithanet.eu/contact). Data stored in IthaGenes are freely available for download in CSV and JSON format.

#### IthaMaps

IthaMaps is a global epidemiology database of haemoglobinopathies, illustrating data on a dynamic global to regional map. For the first release of the IthaMaps database [2], information on relative mutation frequencies were extracted from 89 publications and were illustrated on an interactive map at the country level for a total of 56 countries. Herein, the database and interface have been revamped to allow integration of additional types of information and more detailed data. Specifically, the IthaMaps content can now be divided into three classes: (a) country-specific information on existing haemoglobinopathy-related healthcare policies, (b) country-specific information on the status of major haemoglobinopathies, such as the prevalence, incidence and overall burden of β-thalassaemia and SCD, and (c) relative allele frequencies of specific globin gene mutations at the national and regional level, dynamically linked to corresponding IthaGenes entries.

In addition and through synergies of ITHANET with the HVP GG2020 Challenge, IthaMaps has benefitted significantly from direct contributions to its content from local experts, with 21 scientists annotating or reviewing data for their countries of expertise. In combination with a high volume of extracted information from a total of 287 published reports, IthaMaps organises epidemiological data for 183 countries, thus becoming a valuable tool for stakeholders and policymakers aiming for evidence-based decision making. Table 1 shows the distribution of annotated countries for parameters that describe existing haemoglobinopathy-related healthcare policies, and Table 2 shows the number of annotated countries per continent for epidemiological parameters that describe the status and burden of major haemoglobinopathies.

**Table 1:** The distribution of annotated countries in IthaMaps for each parameter describing existing haemoglobinopathy-related healthcare policies.

| **Healthcare policy** | **Number of countries** |  |  |  |
| --- | --- | --- | --- | --- |
|  | *Yes (National)* | *Yes (Regional)* | *No* | **Total** |
| Prevention programme | 28 | 9 | 37 | **74** |
| SCD Newborn screening | 27 | 8 | 74 | **109** |
| Antenatal screening | 14 | 4 | 14 | **32** |
| Prenatal screening | 21 | 13 | 31 | **65** |
| Haemoglobinopathies patient registry | 20 | 5 | 15 | **50** |
| Rare disease patient registry | 7 | 1 | 15 | **23** |
| Dedicated treatment centres | 33 | 19 | 6 | **58** |
| Blood transfusion availability | 41 | 16 | 2 | **59** |
| Iron chelation availability | 26 | 7 | 2 | **35** |
| MRI facilities | 17 | 9 | 8 | **34** |
| Patient associations | 63 | 1 | 2 | **66** |
| Genetic counselling | 29 | 6 | 4 | **39** |

**Table 2:** The distribution of annotated countries in IthaMaps per continent for each parameter describing the status of major haemoglobinopathies.

|  | Number of countries |  |  |  |  |  | Total |
| --- | --- | --- | --- | --- | --- | --- | --- |
|  | Africa | Asia | Europe | North America | South America | Oceania |  |
| Prevalence of β-thalassaemia carriers | 12 | 43 | 31 | 11 | 6 | 2 | 105 |
| Prevalence of SCD carriers | 30 | 18 | 21 | 32 | 10 | – | 111 |
| Prevalence of α-thalassaemia carriers | 7 | 27 | 3 | – | 2 | – | 39 |
| Prevalence of Hb E carriers | – | 20 | 14 | 2 | 1 | 1 | 38 |
| Prevalence of Hb C carriers | 12 | 2 | 14 | 31 | 6 | – | 65 |
| Expected incidence of β-thalassaemia | 6 | 41 | 28 | 9 | 4 | 1 | 89 |
| Expected incidence of SCD | 7 | 14 | 2 | 10 | 4 | – | 37 |
| Incidence of β-thalassaemia | – | 4 | 4 | – | 1 | – | 9 |
| Incidence of SCD | 51 | 18 | 20 | 25 | 9 | 1 | 124 |
| Incidence of α-thalassaemia | 1 | – | – | – | – | – | 1 |
| Known β-thalassaemia patients | 4 | 32 | 17 | 3 | – | 1 | 57 |
| Known SCD patients | 3 | 16 | 16 | 4 | 1 | – | 40 |

In addition, IthaMaps organises information on relative mutation frequencies, at national or regional level, for 66 countries and for a total of 333 globin gene mutations. To facilitate the identification of the most suitable report, IthaMaps provides detailed information about the parameters of each study, including the sample description (e.g. sample size, population description, specific ethnic group), country region/province, study period and related publication, whilst visualisations of mutation frequencies can be filtered based on the above study parameters.

All epidemiological parameters stored in IthaMaps can be visualised on a worldwide or country-specific interactive map, which can be exported in a variety of image formats. Figure 3 shows three examples of possible illustrations available in IthaMaps: (A) existing healthcare policies with regards to newborn screening for SCD, (B) the worldwide distribution of β-thalassaemia carriers, and (C) the distribution of a selected mutation (e.g. IVS I-5 G>C) in a country (e.g. India) with different frequencies for each province.

**Figure 3:**
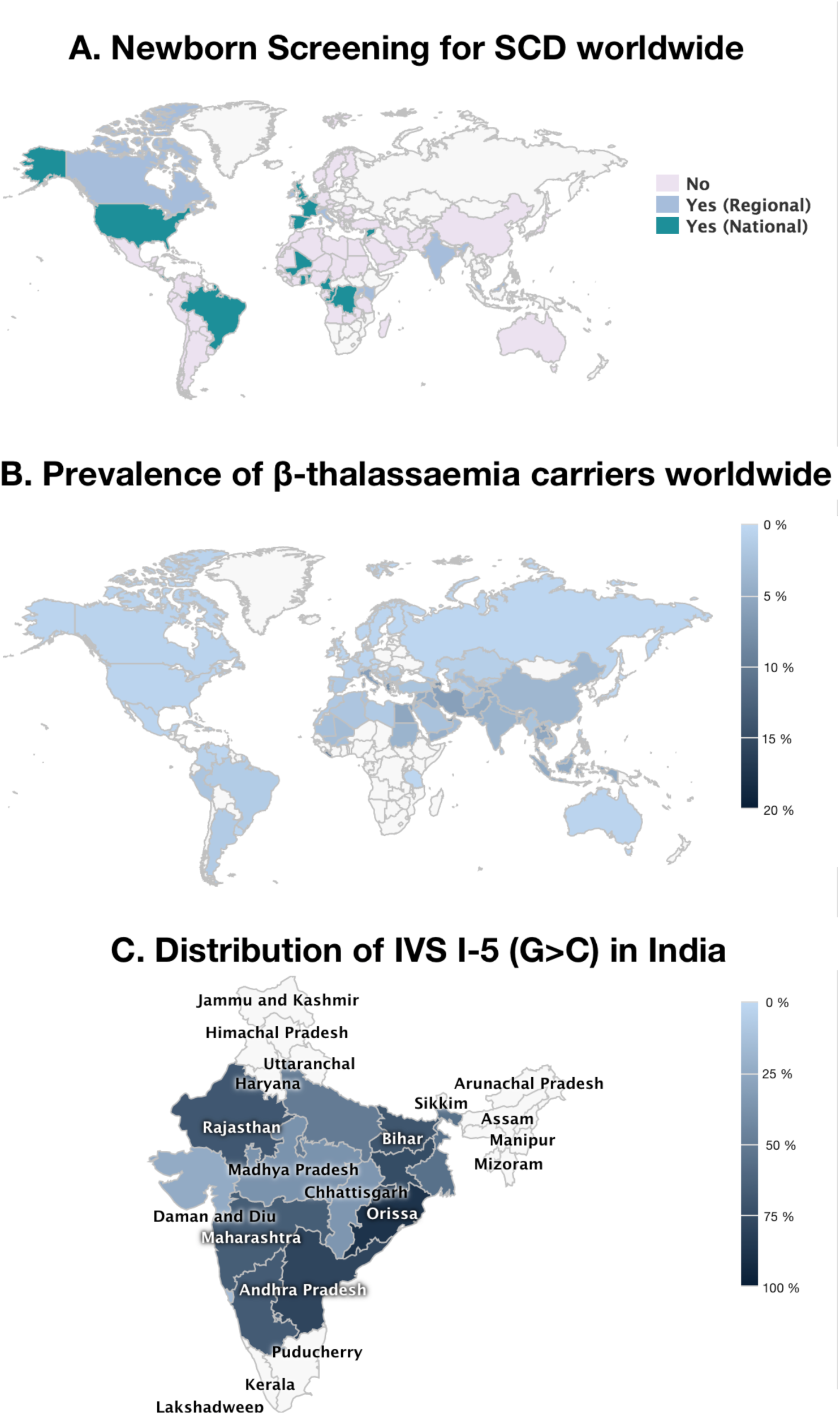
Examples of possible dynamic illustrations available in IthaMaps: (A) existing healthcare policies with regards to prenatal screening, (B) the worldwide distribution of β-thalassaemia carriers, and (C) the relative frequency distribution of mutation IVS I-5 G>C (HBB:c.92+5G>C) in India’s provinces.

#### IthaChrom

Structural variants of the polypeptide globin chains underlying haemoglobinopathies are most commonly detected with the utilisation of high performance liquid chromatography (HPLC). However, their definitive diagnosis is becoming a challenging task, owing to the large and increasing number of newly identified Hb variants, with around 1360 Hb variants currently stored in IthaGenes. IthaChrom aims at supporting the diagnosis of structural haemoglobinopathies by providing digitised anonymised reports (as kindly provided by Bio-Rad Laboratories Inc.) of standard diagnostic HPLC analyses. The interface allows database searches of key data, such as the retention time, specific globin genes, Hb variants, HPLC instruments and methodologies. To date, more than 600 HPLC diagnostic reports are available in IthaChrom and we aim at expanding this section by increasing the catalogue of HPLC methodologies and programmes and by allowing contributions from additional laboratories

## DISCUSSION

The ITHANET Portal is a growing public resource on haemoglobinopathies and, as an interactive community tool, it invites contributions to its content, including news, events, publications and clinical trials, and contact information for clinical, research and diagnostic centres and patient organisations for haemoglobinopathies as well as epidemiological, genetic and diagnostic data. Through these contributions and centralised development, the ITHANET portal aims at expanding its current content with useful tools for scientists and policymakers involved in researching, diagnosing, preventing and treating haemoglobinopathies. Specifically, future developments of the ITHANET portal include the annotation of IthaGenes entries according to the guidelines by the ACMG [31], as part of the ITHANET-GG2020 Expert Panel, the development of epidemiological tools to further facilitate evidence-based policymaking and the implementation of a complete genotype-phenotype database for haemoglobinopathies. Notably, ITHANET is maintained by permanent funding, thus ensuring long-term development and update of the portal, and is an active partner in and beneficiary of primary research data from several projects on haemoglobinopathies. In addition, ITHANET is an active member of the European Reference Network EuroBloodNet for rare haematological diseases and is involved in the development of patient registries for the non-malignant section of the network. Most importantly, through its partnership with the global HVP GG2020 Challenge, ITHANET has become a comprehensive resource for all information relating to the haemoglobinopathies and is evolving into an indispensable tool for the research, prevention and diagnosis of haemoglobinopathies.

## ACKNOWLEDGEMENTS

We are grateful to the following scientists that contributed or curated data: Aurelio Maggio, Bertha Ibarra, Casten W Lederer, Catherine Badens, Celeste Bento, Coralea Stephanou, Cornelis L Harteveld, Esri Voskaridou, Fahd Al-Mulla, Hafizur Rahman, Hassan Syahzuwan, Jacques Elion, Joanne Traeger-Synodinos, Joan-LLuis Vives Corrons, John Old, Leticia Ribeiro, Léon Tshilolo, Marina Kleanthous, Mas Rina Wati Abdoul Hamid, Michael Angastiniotis, Abiageli Nnodu, Paolo Moi, Pavlos Fanis, Petros Kountouris, Salah Al-Humood, Valeriya Kaleva, Xiangmin Xu, Yvonne Daniel, and Bin Alwi Zilfalil. An up-to-date list of ITHANET contributors and curators is provided at http://www.ithanet.eu/community/ithanet-contributors. The authors would like to thank the Cyprus Institute of Neurology and Genetics for funding and computer equipment and for hosting ITHANET. The authors are also grateful to Marco Flamini and Bio-Rad Laboratories, Inc. for providing the HPLC images and reports.

## REFERENCES

1. Modell B, Darlison M. Global epidemiology of haemoglobin disorders and derived service indicators. Bull World Health Organ. 2008;86: 480–487.

2. Kountouris P, Lederer CW, Fanis P, Feleki X, Old J, Kleanthous M. IthaGenes: An Interactive Database for Haemoglobin Variations and Epidemiology. PLoS ONE. Public Library of Science; 2014;9: e103020. doi:10.1371/journal.pone.0103020

3. Cappellini M-D, Cohen A, Eleftheriou A, Piga A, Porter J, Taher A. Guidelines for the clinical management of thalassaemia. 2nd ed. Nicosia: Thalassaemia International Federation; 2008. pp. 1–202.

4. Angastiniotis M, Modell B. Global epidemiology of hemoglobin disorders. Ann N Y Acad Sci. 1998;850: 251–269.

5. Kountouris P, Kousiappa I, Papasavva T, Christopoulos G, Pavlou E, Petrou M, et al. The molecular spectrum and distribution of haemoglobinopathies in Cyprus: a 20-year retrospective study. Sci Rep. 2016;6: 26371. doi:10.1038/srep26371

6. Thein SL. Genetic modifiers of beta-thalassemia. Haematologica. 2005;90: 649–660.

7. Henderson S, Timbs A, McCarthy J, Gallienne A, Van Mourik M, Masters G, et al. Incidence of haemoglobinopathies in various populations-the impact of immigration. Clin Biochem. 2009;42: 1745–1756. doi:10.1016/j.clinbiochem.2009.05.012

8. Kyrri AR, Kalogerou E, Loizidou D, Ioannou C, Makariou C, Kythreotis L, et al. The changing epidemiology of β-thalassemia in the Greek-Cypriot population. Hemoglobin. 2013;37: 435–443. doi:10.3109/03630269.2013.801851

9. Giordano PC, Harteveld CL, Bakker E. Genetic epidemiology and preventive healthcare in multiethnic societies: the hemoglobinopathies. Int J Environ Res Public Health. 2014;11: 6136–6146. doi:10.3390/ijerph110606136

10. Darlison MW, Modell B. Sickle-cell disorders: limits of descriptive epidemiology. Lancet. 2013;381: 98–99. doi:10.1016/S0140-6736(12)61817-0

11. Sherry ST, Ward MH, Kholodov M, Baker J, Phan L, Smigielski EM, et al. dbSNP: the NCBI database of genetic variation. Nucleic Acids Res. 2001;29: 308–311. doi:10.1093/nar/29.1.308

12. Hamosh A, Scott AF, Amberger J, Valle D, McKusick VA. Online Mendelian Inheritance in Man (OMIM). Hum Mutat. 2000;15: 57–61. doi:10.1002/(SICI)1098-1004(200001)15:1<57::AID-HUMU12>3.0.CO;2-G

13. Landrum MJ, Lee JM, Riley GR, Jang W, Rubinstein WS, Church DM, et al. ClinVar: public archive of relationships among sequence variation and human phenotype. Nucleic Acids Res. 2014;42: D980–5. doi:10.1093/nar/gkt1113

14. Cooper DN, Ball EV, Krawczak M. The human gene mutation database. Nucleic Acids Res. 1998;26: 285–287. doi:10.1093/nar/26.1.285

15. Johnston JJ, Biesecker LG. Databases of genomic variation and phenotypes: existing resources and future needs. Human Molecular Genetics. 2013;22: R27–31. doi:10.1093/hmg/ddt384

16. Hardison RC, Chui DHK, Giardine B, Riemer C, Patrinos GP, Anagnou N, et al. HbVar: A relational database of human hemoglobin variants and thalassemia mutations at the globin gene server. Hum Mutat. 2002;19: 225–233. doi:10.1002/humu.10044

17. Fokkema IFAC, Taschner PEM, Schaafsma GCP, Celli J, Laros JFJ, Dunnen den JT. LOVD v.2.0: the next generation in gene variant databases. Hum Mutat. 2011;32: 557–563. doi:10.1002/humu.21438

18. Giardine B, Borg J, Higgs DR, Peterson KR, Philipsen S, Maglott D, et al. Systematic documentation and analysis of human genetic variation in hemoglobinopathies using the microattribution approach. Nature Genetics. 2011;43: 295–301. doi:10.1038/ng.785

19. Lederer CW, Basak AN, Aydinok Y, Christou S, El-Beshlawy A, Eleftheriou A, et al. An electronic infrastructure for research and treatment of the thalassemias and other hemoglobinopathies: the Euro-mediterranean ITHANET project. Hemoglobin. 2009;33: 163–176. doi:10.1080/03630260903089177

20. Robinson HM. Increasing the involvement of diverse populations in genomics-based health care-lessons from haemoglobinopathies. Journal of community genetics. Springer Berlin Heidelberg; 2017;43: 295–8. doi:10.1007/s12687-017-0327-3

21. Huisman THJ, Carver MFH, Efremov GD. A Syllabus of Human Hemoglobin Variants. 2nd ed. Augusta, GA, USA: The Sickle Cell Anemia Foundation; 1996.

22. Huisman THJ, Carver MFH, Baysal E. A Syllabus of Thalassemia Mutations. Augusta, GA, USA: The Sickle Cell Anemia Foundation; 1997.

23. Steinberg MH, Forget BG, Higgs DR, Weatherall DJ. Disorders of Hemoglobin. 2nd ed. Cambridge University Press; 2009.

24. Old J, Angastiniotis M, Eleftheriou A, Galanello R, Harteveld CL, Petrou M, et al. Prevention of Thalassaemias and Other Haemoglobinopathies. 2nd ed. Nicosia, Cyprus: Thalassaemia International Federation; 2013.

25. Thein SL. The molecular basis of β-thalassemia. Cold Spring Harbor Perspectives in Medicine. 2013;3: a011700. doi:10.1101/cshperspect.a011700

26. Higgs DR. The molecular basis of α-thalassemia. Cold Spring Harbor Perspectives in Medicine. 2013;3: a011718. doi:10.1101/cshperspect.a011718

27. Mottaz A, David FPA, Veuthey A-L, Yip YL. Easy retrieval of single amino-acid polymorphisms and phenotype information using SwissVar. Bioinformatics. 2010;26: 851–852. doi:10.1093/bioinformatics/btq028

28. Danjou F, Anni F, Perseu L, Satta S, Dessì C, Lai ME, et al. Genetic modifiers of β-thalassemia and clinical severity as assessed by age at first transfusion. Haematologica. Haematologica; 2012;97: 989–993. doi:10.3324/haematol.2011.053504

29. Robinson PN, Köhler S, Bauer S, Seelow D, Horn D, Mundlos S. The Human Phenotype Ontology: A Tool for Annotating and Analyzing Human Hereditary Disease. The American Journal of Human Genetics. Elsevier; 2008;83: 610–615. doi:10.1016/j.ajhg.2008.09.017

30. Köhler S, Doelken SC, Mungall CJ, Bauer S, Firth HV, Bailleul-Forestier I, et al. The Human Phenotype Ontology project: linking molecular biology and disease through phenotype data. Nucleic Acids Res. 2014;42: D966–74. doi:10.1093/nar/gkt1026

31. Richards S, Aziz N, Bale S, Bick D, Das S, Gastier-Foster J, et al. Standards and guidelines for the interpretation of sequence variants: a joint consensus recommendation of the American College of Medical Genetics and Genomics and the Association for Molecular Pathology. Genet Med. Nature Publishing Group; 2015. doi:10.1038/gim.2015.30

